# E-cigarettes increase the risk of adenoma formation in murine colorectal cancer model

**DOI:** 10.1101/2024.08.23.609469

**Authors:** Ibrahim M Sayed, Anirban Chakraborty, Kaili Inouye, Leanne Dugan, Stefania Tocci, Ira Advani, Kenneth Park, Tapas K Hazra, Soumita Das, Laura E. Crotty Alexander

**Author notes:** Co-corresponding authors: Ibrahim M Sayed: Department of Biomedical & Nutritional Sciences, Zuckerberg College of Health Sciences, University of Massachusetts Lowell, Lowell, MA 01854, USA., Soumita Das: Department of Biomedical & Nutritional Sciences, Zuckerberg College of Health Sciences, University of Massachusetts Lowell, Lowell, MA 01854, USA., Laure E. Crotty Alexander: Department of Medicine, University of California, San Diego, CA, 92093, USA.

## Abstract

**Background:** E-cigarettes (E.cigs) cause inflammation and damage to human organs, including the lungs and heart. In the gut, E.cig vaping promotes inflammation and gut leakiness. Further, E.cig vaping increases tumorigenesis in oral and lung epithelial cells by inducing mutations and suppressing host DNA repair enzymes. It is well known that cigarette (cig) smoking increases the risk of colorectal cancer (CRC). To date, it is unknown whether E.cig vaping impacts CRC development.

**Methods:** A mouse model of human familial adenomatous polyposis (CPC-APC) was utilized wherein a mutation in the adenomatous polyposis coli (APC) gene, CDX2-Cre-APC^Min/+^, leads to the development of colon adenomas within 16 weeks. Mice were exposed to air (controls), E.cig vaping, cig, or both (dual exposure). After 4 weeks of 2-hour exposures per day (1 hour of each for dual exposures), the colon was collected and assessed for polyp number and pathology scores by microscopy. Expression of inflammatory cytokines and cancer stem cell markers were quantified. DNA damage such as double-strand DNA breaks was evaluated by immunofluorescence, western blot and gene-specific long amplicon qPCR. DNA repair enzyme levels (NEIL-2, NEIL-1, NTH1, and OGG1) were quantified by western blot. Proliferation markers were assessed by RT-qPCR and ELISA.

**Results:** CPC-APC mice exposed to E.cig, cig, and dual exposure developed a higher number of polyps compared to controls. Inflammatory proteins, DNA damage, and cancer stemness markers were higher in E-cig, cig, and dual-exposed mice as well. DNA damage was found to be associated with the suppression of DNA glycosylases, particularly with NEIL-2 and NTH1. E.cig and dual exposure both stimulated cancer cell stem markers (*CD44, Lgr-5, DCLK1, and Ki67).* The effect of E.cigs on polyp formation and CRC development was less than that of cigs, while dual exposure was more tumorigenic than either of the inhalants alone.

**Conclusion:** E.cig vaping promotes CRC by stimulating inflammatory pathways, mediating DNA damage, and upregulating transcription of cancer stem cell markers. Critically, combining E.cig vaping with cig smoking leads to higher levels of tumorigenesis. Thus, while the chemical composition of these two inhalants, E.cigs and cigs, is highly disparate, they both drive the development of cancer and when combined, a highly common pattern of use, they can have additive or synergistic effects.

## Introduction

E-cigarettes (E.cigs) are a non-combustible form of tobacco and are especially popular among youth. E-cigs were first introduced in the mid-2000s, and have become widespread in the US, especially among adolescents. Approximately 10 million US adults and 2 million kids are active E.cig users [1–4]. E.cig liquids (e-liquids) consist of vehicle solvents such as propylene glycol (PG) and glycerin, commonly called vegetable glycerin (VG), combined with nicotine, which are heated and aerosolized within e-devices. A large variety of flavoring chemicals (∼7,000) are added to e-liquids to increase appeal [5]. It is thought that switching from conventional tobacco to E.cigs can bypass the harmful effects of conventional smoking [6], however, there are high rates of dual use of both cigarettes and E.cigs.

Vaping has been associated with some adverse health effects, including lung injury (EVALI) [7,8]. Chronic inhalation of E.cig affects the inflammatory states in the lungs and circulation and can increase the risk of infectious diseases [9]. Our recent study showed that E.cig vaping disrupts gut barrier integrity and induces inflammation and gut leakiness [10]. Gut leakiness and intestinal barrier defects with microbial dysbiosis are risk factors for colorectal carcinoma (CRC)[11]. It is well established that cigarette smoking increases the risk of CRC, due to smoking-induced DNA damage in the colon [12–14]. While cigarette smoking has been clearly linked to the development of CRC [15–17], there is no data directly linking E.cig vaping with CRC development.

CRC is the second leading cause of death in the USA [18]. Several factors can contribute to the development of CRC such as genetic mutations, environmental factors, lifestyles, and microbial dysbiosis [19]. CRCs are primarily caused by the loss of tumor-suppressor genes, such as APC [20], and almost 80% of CRC cases are due to APC mutations [21]. The loss of APC gene is the cause of familial adenomatous polyposis (FAP) [22]. CPC-APC (CDX2-Cre-APCmin with a mutation in the APC gene) mice are a mouse model of FAP and CRC that spontaneously develop multiple colonic polyps (adenomas) and rectal bleeding at around 16 weeks [23]. Cigarette smoke exposure increases the development of adenomas in APC^Min/+^ mice by altering intestinal permeability [24].

Toxicological studies have reported the presence of heavy metals, formaldehyde and acrolein in E.cig aerosols, commonly called vapor, raising health concerns [25]. In addition, PG, VG, flavorings, and contaminants can be toxic [26,27]. E.cig vapor has been shown to induce DNA damage and accumulative mutations [28,29]. E.cig effects were comparable to or slightly higher than the effect mediated by mainstream smoke extracts, suggesting potential carcinogenic effects of E.cig components [28]. Using 3D human and murine organoids and 2D enteroid-derived monolayers, we have shown that exposure to E.cig vapor increased the bacterial translocation inside the gut, upregulated inflammatory responses, and altered DNA damage responses [10].

Following DNA damage such as oxidation, deamination, and alkylation, the host base excision repair (BER) proteins/ DNA glycosylases can excise the damaged based and the downstream steps correct the DNA damage to prevent accumulation of the DNA mutation and development of CRC [30,31]. Suppression and downregulation of DNA glycosylases, especially NEIL-2 and NEIL-1, have been associated with the development of lung cancer [32]. Our studies showed that the gastric cancer-associated microbes *Helicobacter pylori* and CRC-associated microbe *Fusobacterium nucleatum* downregulate epithelial DNA glycosylase NEIL-2, increase inflammation and leads to the accumulation of DNA damage and cancers [33,34]. Exposure to E.cig vapor causes a suppression of OGG1 in oral and lung epithelial cells resulting in the accumulation of reactive oxygen species, 8-hydroxy-2’-deoxyguanosine (8-oxo-dG) bases which are genotoxic [28]. However, the effect of E.cig exposure on other DNA glycosylases and in the colon has not been studied.

In this study, we assessed the impact of E.cig vapor inhalation alone and in combination with cigarette smoke on the development and progression of CRC. We exposed CPC-APC mice to E.cig vapor, cigarette smoke, and both inhalants (dual exposure), or air (control) and assessed CRC development via tumor number, tumor size, inflammation, DNA damage, and intestinal stemness markers. We also assessed the damaging effects of E.cigs on genomic DNA and on host DNA repair enzymes.

## Methods

### 1 ) Animals and Inhalant Exposures

CPC-APC (CDX2-Cre-APCmi) mice were housed, bred, enrolled in the experimental design, euthanized according to the University of California San Diego Institutional Animal Care and Use Committee (IACUC) policies under animal protocol numbers (s18086 and s16021). Prior to randomization at the beginning of all studies, all mice of the same sex were group housed in a large sterile container for 2 hours, to allow free mixing of microbiota amongst individuals.

CPC-APC mice of both genders, aged 9-12 weeks, were exposed for 2 hours daily to inhalants for 4 weeks via whole-body exposure chambers (inExpose system by SciReq). All mice were placed into the chambers for 2 hours per day. CPC-APC mice were divided into four groups: a) Control, air-exposed, b) cig smoke-exposed (positive control), c) E.cig aerosol, commonly called E.cig vapor (EV), exposed, and d) Dual exposure to both cig and E.cig, to seek if co-use of both increases risk of CRC (Figure 1A). Each exposure type had a dedicated chamber, tubing and pump, to prevent cross-contamination. For EV generation, the following vaping parameters were used: inhalation cycles per minute (Nc = 3), drag time (Dt = 4s), peak inhalation flow (Qi = 2L/s), and electrical driving power (W = 6-8 watts based on the JUUL coil of 1.6 ohms and voltage of 3.7), using e-liquid composition of 30% propylene glycol (PG), 70% glycerin (Gly) and 59 mg/mL nicotine, which match the most popular e-liquids used by 4th generation, pod-based e-devices (JUUL, FLUM, Sourin, etc.). Aliquots of each batch of e-liquid were stored at −80°C for evaluation by Mass spectrometry (UCSD CoRE facility). For cigarette smoke generation, Nc = 1, Dt = 1s, and Qi 2L/s were used, and 1R6F research cigarettes with the filter removed were replaced when <1cm remains (∼1 cigarette every 8-10 minutes). For Dual exposures, mice were exposed to EV for 60 min and Cig for 60 min each day, yielding 2-hours of inhalant exposure daily. Mice were exposed for 4 weeks, colons of mice were harvested, and the polyp number and area were measured.

**Figure 1:**
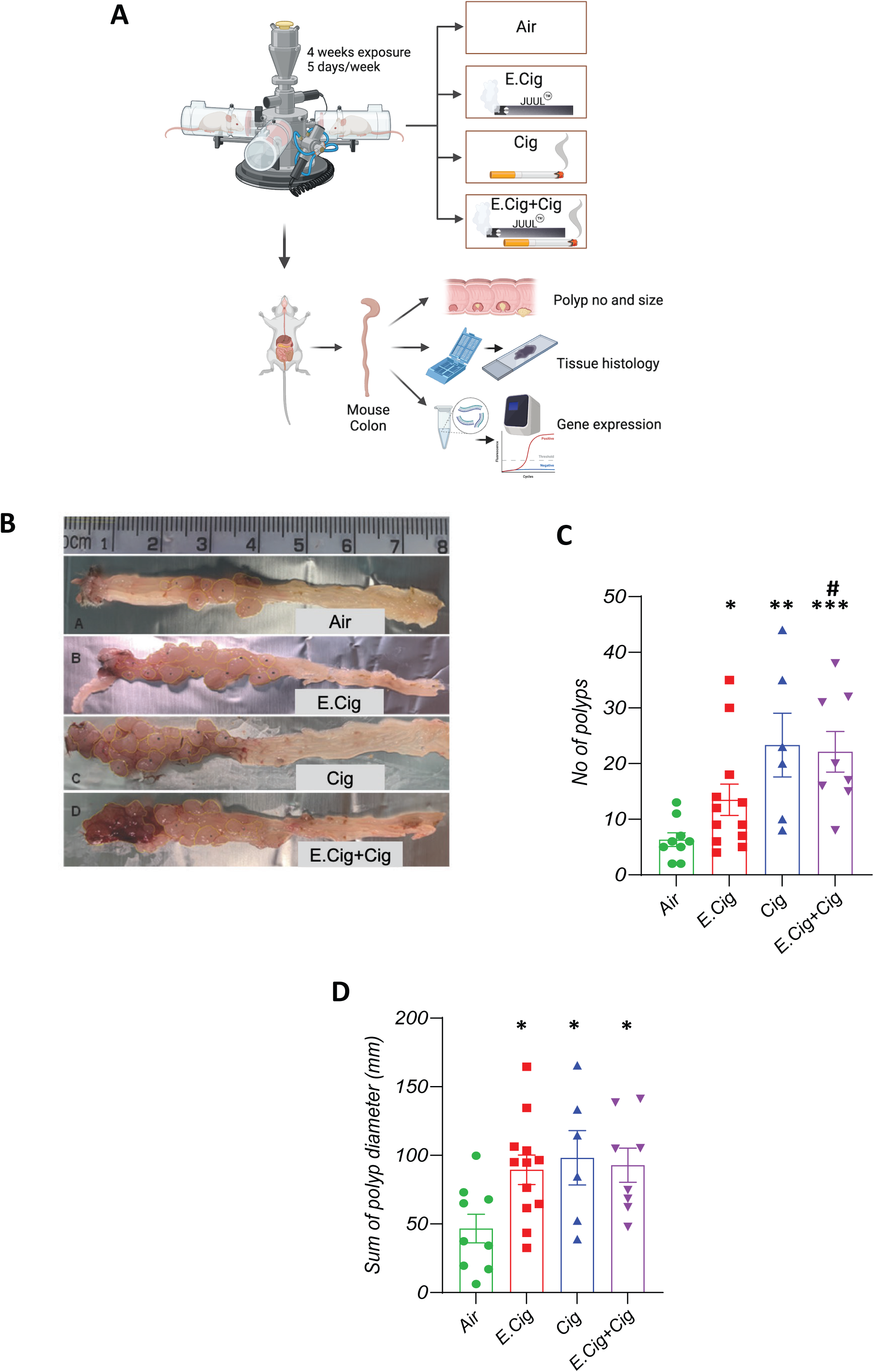
E.cig and Dual exposures increase polyp formation in CRC-APC mice. CPC-APC mice (murine CRC model) were exposed to Air, E.cig, Cig, and E.cig + Cig (Dual exposure). Colonic tissue was collected after 4 weeks of exposure. (A) Experimental design infographic. (B) Representative mouse colon pictures from each exposure. (C) The number of polyps per mouse (n = 6-12 per group). (D) The sum of polyp diameter (mm) was calculated per mouse (n=6-12 per group). Each dot represents one mouse. Data shown are means ± SEM. \**P*-value < 0.05, \*\**P*-value < 0.01, and \*\*\**P*-value < 0.001, as determined by the Mann-Whitney test. * indicates comparison to air-exposed controls; # indicates comparison to E.cig exposed mice.

### 2) Quantification of Colon Adenomas

Upon harvesting, each colon was cleaned and laid flat next to a ruler. Pictures of the colon were taken. Then, the pictures were uploaded into ImageJ software, which was used to count the number of polyps present in the mouse colons after exposure. The area of each polyp was also determined, and the combined total area of all polyps was calculated.

### 3) Histologic Analysis of the Colon

Colon tissues were fixed using 10 % formalin overnight. Tissues were embedded in paraffin blocks, sectioned onto slides, and stained with H&E. The histologic score was determined by assessment of mucosal architecture and crypt loss and infiltration of immune cells in different colon regions (lamina propria, mucosa, and submucosa) as described previously [35].

### 4) Inflammatory and Proliferation Gene Expression Assessment by RT-qPCR

Total RNA was extracted from polyp regions (less than 25 mg) using the ZymoRNA miniprep kit (Zymo Research) and complementary DNA (cDNA) was reverse transcribed using qScript cDNA SuperMix (Quantabio). Real-time RT-PCR was performed using 2x SYBR Green qPCR Master Mix (Biomake) and Bio-Rad CFX Opus 96 Real-Time PCR System (Bio-Rad). 18srRNA and GAPDH were used as housekeeping genes for normalization. The ΔΔCt method was used for the calculation as done previously [36]. The relative fold change in the transcript level was calculated. The primer sequences used are listed in Table 1.

**Table 1.**
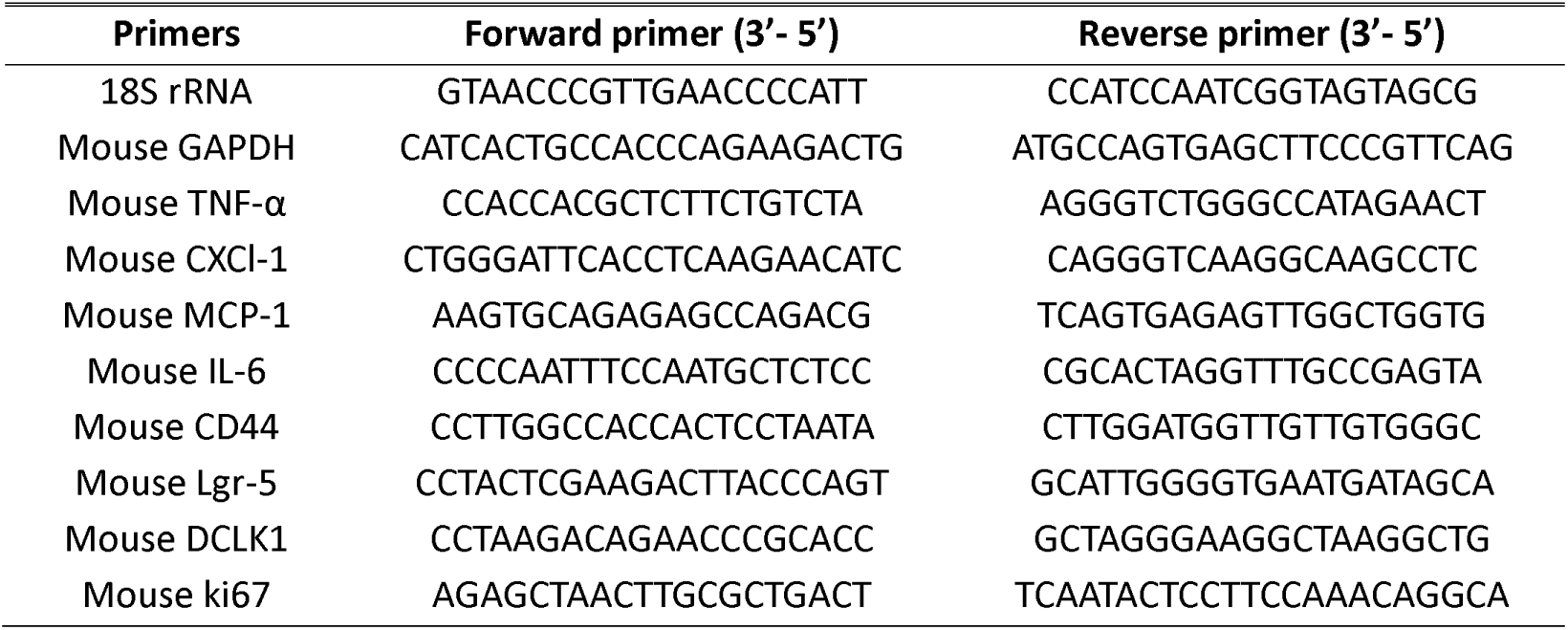
Primer Sequences for RT-qPCR.

### 5) Quantification of DNA Strand Breaks by Long Amplicon (LA) qPCR

The detection of accumulated DNA strand breaks was performed as described previously [33]. Briefly, genomic DNA was extracted from polyps (one polyp per mouse and 3 mice/ each group of exposure) and levels of DNA strand breaks in polß (6.5 Kb) and globin (8.7 Kb) were quantified as described previously [37]. Briefly, the extracted DNA was amplified using the following condition: 94 °C for 30 s (94 °C for 30 s, 55–60 °C for 30 s depending on the oligo annealing temperature, and 65 °C for 10 min) for 25 cycles and 65 °C for 10 min. The sequences of oligo were described before [33,37]. Short amplicons for these genes were used to normalize the amplification of long amplicons.

### 6) Immunofluorescence for Detection of DNA Damage

Colon sections were deparaffinized in xylene, rehydrated in graded ethanol, and rinsed in Tris-phosphate-buffered saline (TBS). Heat-induced antigen retrieval was performed in a microwave for 30 minutes at 98°C in 0.01 M citrate buffer (pH6). After cooling for 20 minutes, sections were incubated in 1% Triton X-100 for 30 minutes, blocked in TBST with 10% donkey serum for 1 hour then incubated with primary rabbit monoclonal antibody Phospho-Histone H2A.X (Ser139), to detect γH2AX, overnight at 4°C (1:250, Cell Signaling, #9718). After subsequent washes in TBST, slides were incubated with secondary Donkey anti-Rabbit IgG (H+L) Alexa fluor plus 488 (1:500, Invitrogen, A32790) for 1 hour in the dark. Immunofluorescence (IF) images were taken with a 40X objective using a Leica SP8 II confocal laser scanning microscope.

### 7) Western Blot Detection of DNA Damage and DNA Repair Proteins

Colon polyps (one polyp per mouse and 3 mice/ each group of exposure) were lysed using the RIPA buffer and the assessment of the level of base excision repair (BER) enzymes was performed as described before [33]. Briefly, after separation of the proteins via acrylamide gels and transfer onto nitrocellulose membranes, membranes were blocked with 5% w/v skimmed milk and incubated with appropriate primary rabbit-derived antibodies (NEIL2, NTH1, and OGG1; in-house antibodies used in 1:500 dilutions). gH2AX and H2AX antibodies were purchased from Cell Signaling Technology (cat# 9718 and 7613, respectively). Histone deacetylase 2 (HDAC2; GeneTex, Irvine, CA, USA) was used as a loading control. After washing steps, the membranes were incubated with anti-isotype secondary antibody (GE Healthcare) conjugated with horseradish peroxidase in 5% skimmed milk at room temperature. All the western blot (WB) images were quantified using the ImageJ automated digitizing system based on three independent gel images (n = 3).

### 8) Quantification of Proliferation

Colonic polyp regions (one polyp per mouse and 5 mice/ each group of exposure) and non-polyp regions were homogenized in phosphate-buffered saline (PBS). The protein concentrations were measured, and equal amounts of each sample (∼10 ug) were used in the Mouse Ki-67 (Ki-67) ELISA Kit (MyBiosource, Catalogue # MBS1601117) according to manufacturer’s instructions. The standard curve was developed using Ki67 standards, and the quantity of Ki67 in each sample was determined by measuring the absorbance at 450 nm.

### 9) Statistics

Analyses were conducted in Graphpad prism V10 (La Jolla, CA). Data is presented as mean ± SEM and analyzed using non-parametric Student’s t-test and One-Way ANOVA as specified. Results are considered significant if the *p*-value is less than 0.05.

## Results

### E.cig and Dual exposures increase adenomatous polyp formation

Previous studies have shown that cigarette smoke (cig) increases the risk of CRC and adenoma formation in the CPC-APC murine model by affecting gut microbiota and gut barrier integrity [24,38]. Further, our previous study showed that E.cig vaping induces inflammation and gut leakiness in healthy gut and wild-type mice [10]. Here we assessed the impact of E.cig vaping and Dual exposure to both E.cig and cig on CRC risk and promotion of adenoma formation using the CPC-APC mouse model [23] (Figure 1A). As expected, the polyp number and area were increased in the CPC-APC mice after cigarette smoke exposure (Figure 1B-1D). Interestingly, we found that E.cig exposure significantly increased the polyp number and area in exposed mice compared to air-exposed controls (Figure 1B-1D). Regarding Dual exposure (E.cig + cig), polyp number and polyp area were also increased (Figure 1B-1D), and the number of polyps were significantly higher in Dual-exposed mice relative to E.cig exposed mice (Figure 1C).

### E.cig and Dual exposures increase inflammatory responses in the colon

Inflammation in the colon and tissue damage are linked to an increase risk of CRC [39,40]. Therefore, we assessed if E.cig exposure promotes adenoma formation by affecting the inflammatory signaling pathways. To this end, colons of exposed mice were stained with H&E (Figure 2A) and the pathology score was calculated (Figure 2B). The pathology score is based on the degree of infiltration of immune cells in the colon and the architecture of colon mucosa [35]. The pathology score was significantly higher in E.cig-exposed mice, cig-exposed mice, and Dual-exposed mice compared to air-exposed controls (Figure 2B). We also assessed transcript levels of inflammatory cytokines TNF-α, IL-6, MCP-1, and CXCL-1 (IL-8 homolog) in the polyps. We selected these cytokines since our previous studies showed that E.cig vaping increased inflammatory transcripts in the gut, particularly MCP-1, IL-6, and IL-8, leading to intestinal barrier defects [10]. Also, our previous study showed that cancer-associated microbes could promote CRC progression in the enteroid-derived monolayer isolated from the colon of APC-min mice by affecting CXCL-1/IL-8 cytokines [33]. Cigarette, E.cig, and Dual exposures significantly increased transcript levels of TNF-α, CXCL-1, MCP-1, and IL-6 in the polyp regions compared to air-exposed controls (Figure 2C-2F). Interestingly, the levels of CXCL-1, MCP-1, and IL-6 and were significantly higher in the polyps of Dual-exposed mice compared to the polyps of E.cig-exposed mice (Figure 2D-2F).

**Figure 2:**
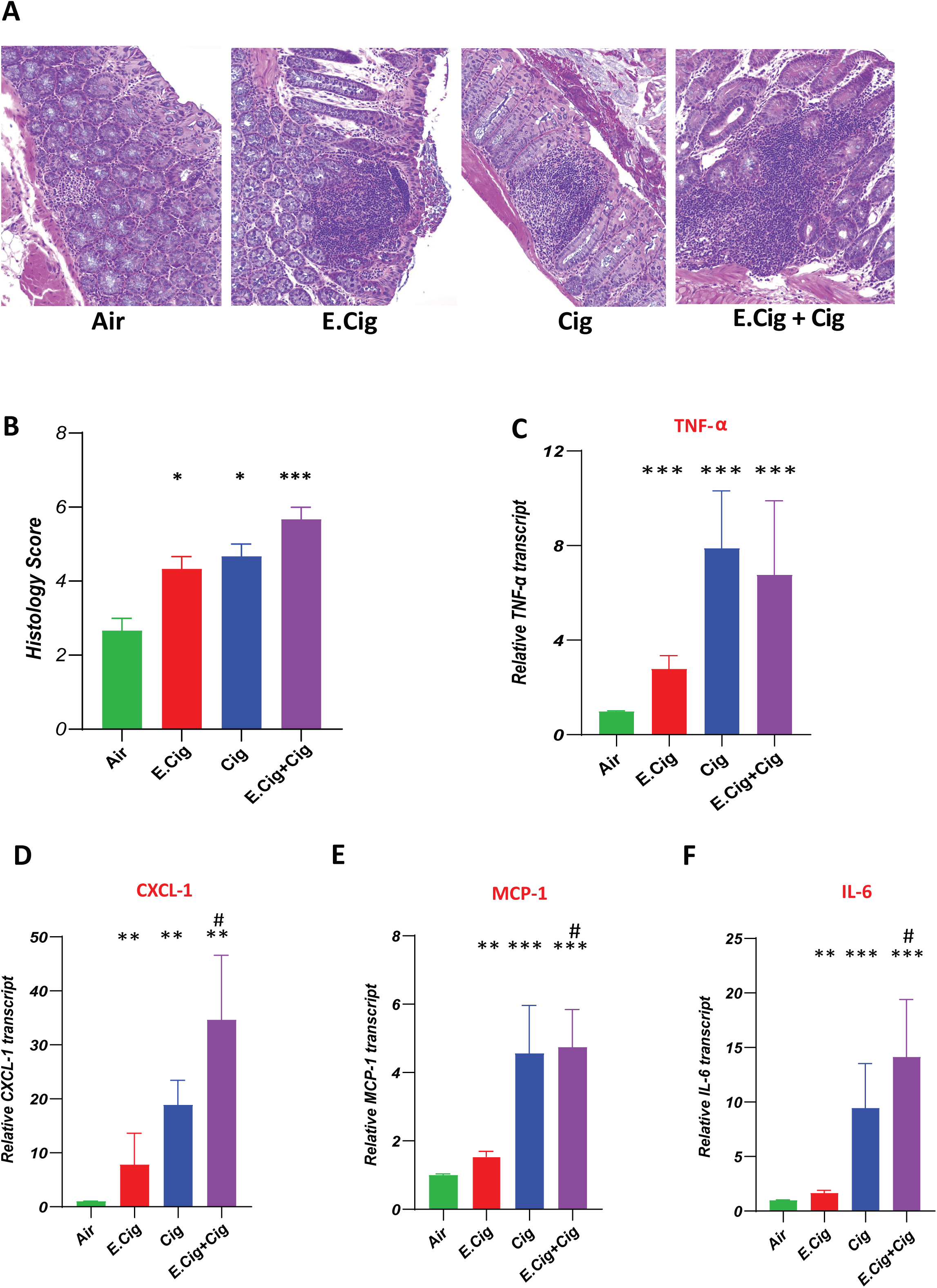
E.cig and Dual exposures increase inflammatory responses in the colon. (A) Representative images of colons of CPC-APC mice from the four different conditions, stained with H&E. (B) Colon histology score was determined based on quantity of inflammatory cells in different regions (lamina propria, mucosa, and submucosa) and the morphology of mucosal architecture. (C-F) mRNA levels of inflammatory cytokines in colonic polyps, including TNF-α (C), CXCL-1 (D), MCP-1 (E), and IL-6 (F). Data presented as means ± SEM. \**P*-value < 0.05, \*\**P*-value < 0.01, and \*\*\**P*-value < 0.001, as determined by the Mann-Whitney test. * indicates comparison to air-exposed controls; # indicates comparison to E.cig exposed mice, P < 0.05.

### E.cig and Dual exposures mediate accumulation of mutations via induction of dsDNA breaks and suppression of host DNA glycosylases

Induction of DNA damage and failure to repair damage leads to the accumulation of mutations and genomic instability which promote cancer initiation and progression [41]. In vivo animals exposed to E.cig had toxic metabolites from nicotine which led to O(6)-methyl-2’-deoxyguanosine DNA adducts which increase mutagenesis in human lung and bladder epithelial cells [42]. E.cig aerosols are genotoxic to human oral and lung epithelial cells by increasing oxidative DNA damage [28]. Therefore, we asked if E.cig exposure promotes adenoma formation in the CRC murine model via inducing dsDNA breaks and mutations. To this end, we assessed the level of γH2AX (a marker of DNA double-strand breaks) in the colon of exposed mice by IF and WB (Figure 3A-3B). Levels of γH2AX were higher in E.cig-exposed mice, cig-exposed mice, and Dual-exposed mice compared to air-exposed controls (Figure 3A-3C). The highest level of γH2AX was seen in Dual-exposed mice (Figure 3A-3C). Then, we tested the effect of exposures on host DNA repair proteins, more specifically DNA glycosylases (NEIL1, NEIL2, NTH, and OGG1). Previous studies have shown that conventional smoking and E.cig mediate host DNA damage by suppressing DNA glycosylase expression [28,42]. We found that E.cig, cig, and Dual exposures reduce expression of all the tested DNA glycosylases (NEIL2, NTH1, and NEIL1) except OGG1 (Figure 3D-3E). Also, we quantified DNA strand breaks in Polβ and β-globin genes from polyps using LA-qPCR (Figure 3F-3G). We observed a significantly higher level of DNA strand-break accumulation in both the Polβ and β-globin genes following exposure to E.cig, cig, and Dual exposures compared to air-exposed controls (Figure 3F-3G). Collectively, E.cig exposure promoted polyp formation in CPC-APC mice by inducing DNA breaks, DNA mutations, and downregulating DNA repair proteins/ DNA glycosylases.

**Figure 3:**
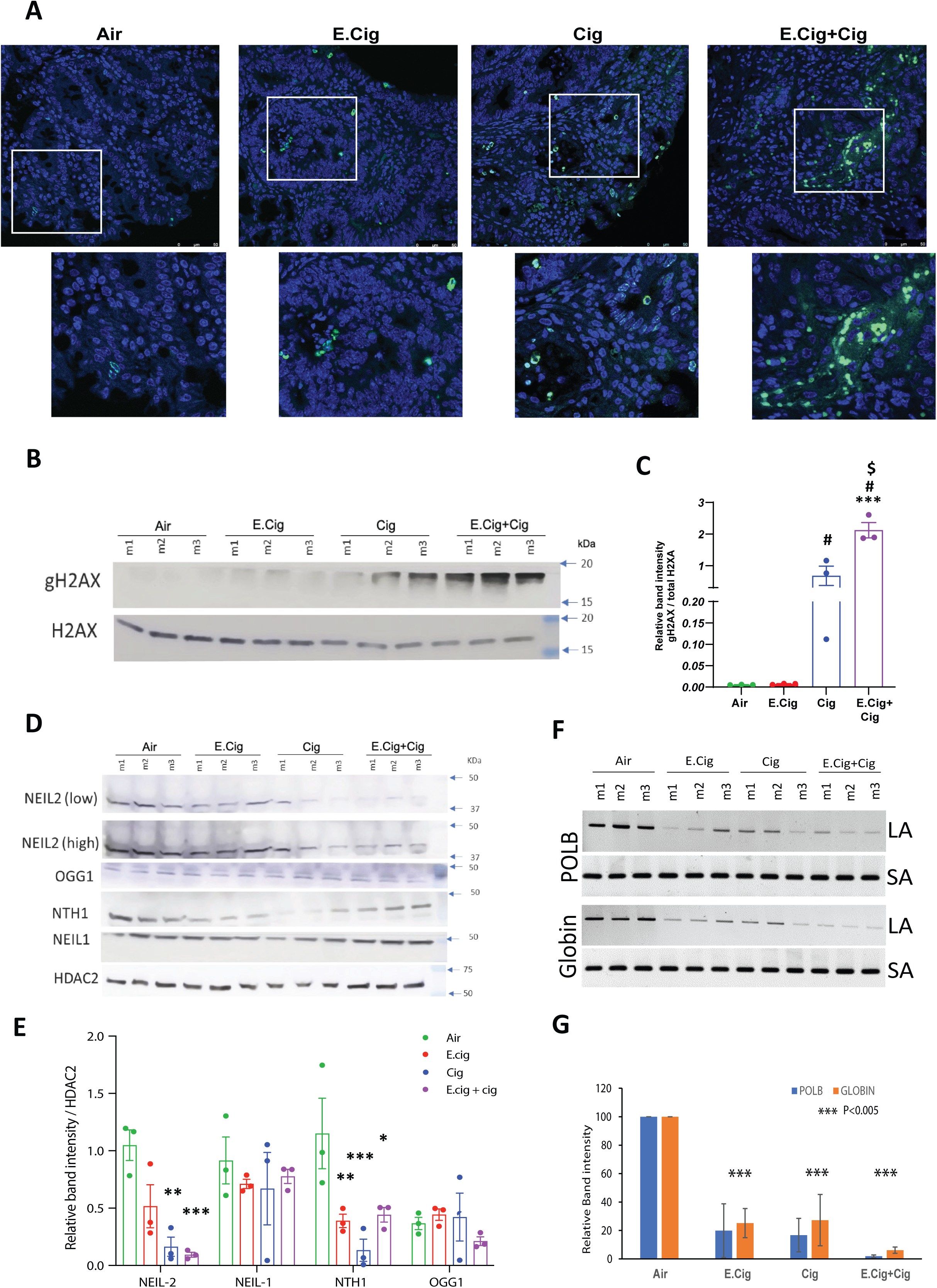
E.cig and Dual exposures induce double-strand (ds) DNA breaks and suppress host DNA repair glycosylases. (A) Colonic polyps were stained for γH2AX for dsDNA breaks (green) and DAPI for nuclei staining (blue)and assessed with immunofluorescence (IF). Images were taken using 40X magnification, and the scale bar is 50 μm. (B) Immunoblot to γH2AX levels in the extract of polyps. Total H2AX was used for normalization. Three independent experiments (n = 3) were performed, and one representative figure is shown. (C) The band density of γH2AX normalized to total H2AX was calculated for each sample (analyzed with one-way ANOVA). (D) Immunoblot of NEIL2, NEIL-1, NTH1, and OGG1 levels in the extract of polyps. HDAC2 (histone deacetylase 2) was used as loading control. Three independent experiments (n = 3) were performed, and one representative figure is shown. (E) The band density of DNA glycosylases (NEIL-1, NEIL-2, NTH1, and OGG1) normalized to HDAC2 was calculated in each sample (analyzed with two-way ANOVA). (F) The polyps of exposed mice were harvested for genomic DNA isolation. Long amplicon quantitative PCR (LA-qPCR) was performed to evaluate the level of DNA strand-break accumulation. Representative gels show the amplification of each long fragment (∼7–8 kB; upper panel) normalized to that of a short fragment (∼250 bp; lower panel) of the corresponding (Pol β and β-Globin) genes. (G) The relative band density in (F) was calculated, and compared with air-exposed controls. Data presented as means ± SEM. \**P*-value < 0.05, \*\**P*-value < 0.01, and \*\*\**P*-value < 0.001. * indicates comparison to air-exposed controls; # indicates comparison to E.cig exposed mice, P < 0.05; $ indicates comparison to cig-exposed mice, P < 0.05.

### E.cig and Dual exposures upregulate cancer stem cell markers and proliferation markers in the colon to initiate the cancer program

Previous studies have shown that cigarette smoke promotes proliferation and epithelial-mesenchymal transition (EMT), and increases the activity of cancer stem cells (CSCs) in the lung [43], kidney [44], bladder [45], skin [46], liver [47], and pancreas [48]. The CSCs are a subpopulation of tumor cells that promote cancer initiation, progression, metastasis, and resistance to chemotherapy and radiation [49]. There are several markers for CSCs such as cluster of differentiation 44 (CD44) [50], Leucine-rich repeat-containing G-protein-coupled receptor 5 (LGR5) [51], and Doublecortin-like kinase 1 (Dclk1) [52]. Ki-67 is another marker that plays a crucial role in the maintenance of colon CSCs, and it is a marker of cell proliferation [53]. We focused on these four markers since previous studies showed that these markers were associated with adenoma formation and CRC in the APC mouse model [52,54–56]. We queried whether E.cig exposure affects the level of colon CSCs. To this end, we analyzed the transcript levels of *CD44, Lgr-5, DCLK1, and Ki67* in the polyp regions across exposure groups (Figure 4A). Cigarette smoke significantly upregulated the four CSC markers (∼4-8 fold) compared to air-exposed controls (Figure 4B-E). Likewise, E.cig induced expression of *CD44, Lgr-5, and Ki67*, but not *DCLK1*, in polyp regions (Figure 4B-E). Dual exposure caused a significant stimulation in the transcriptome of the CSCs markers (Figure 4B-E), and mRNA levels of *CD44* and *DCLK1* were significantly higher in Dual-exposed mice compared to E.cig-exposed mice (Figure 4B-C). To confirm our findings, we assessed Ki67^+^ cells in polyp regions by measuring protein levels of Ki67 in whole cell extracts by ELISA (Figure 4F). First, we assessed the the Ki67 in the non-involved region and polyp region from the same mouse,. As expected, the level of Ki67 was significantly higher in the polyp regions compared to the non-involved regions as it is a marker of cell proliferation (Figure 4F). Then we compared the Ki67 protein level in all exposure groups of mice in the polyp region. We found that the level of Ki67 was significantly higher in E-cig, cig, and Dual-exposed mice compared to air-exposed controls (Figure 4G).

**Figure 4:**
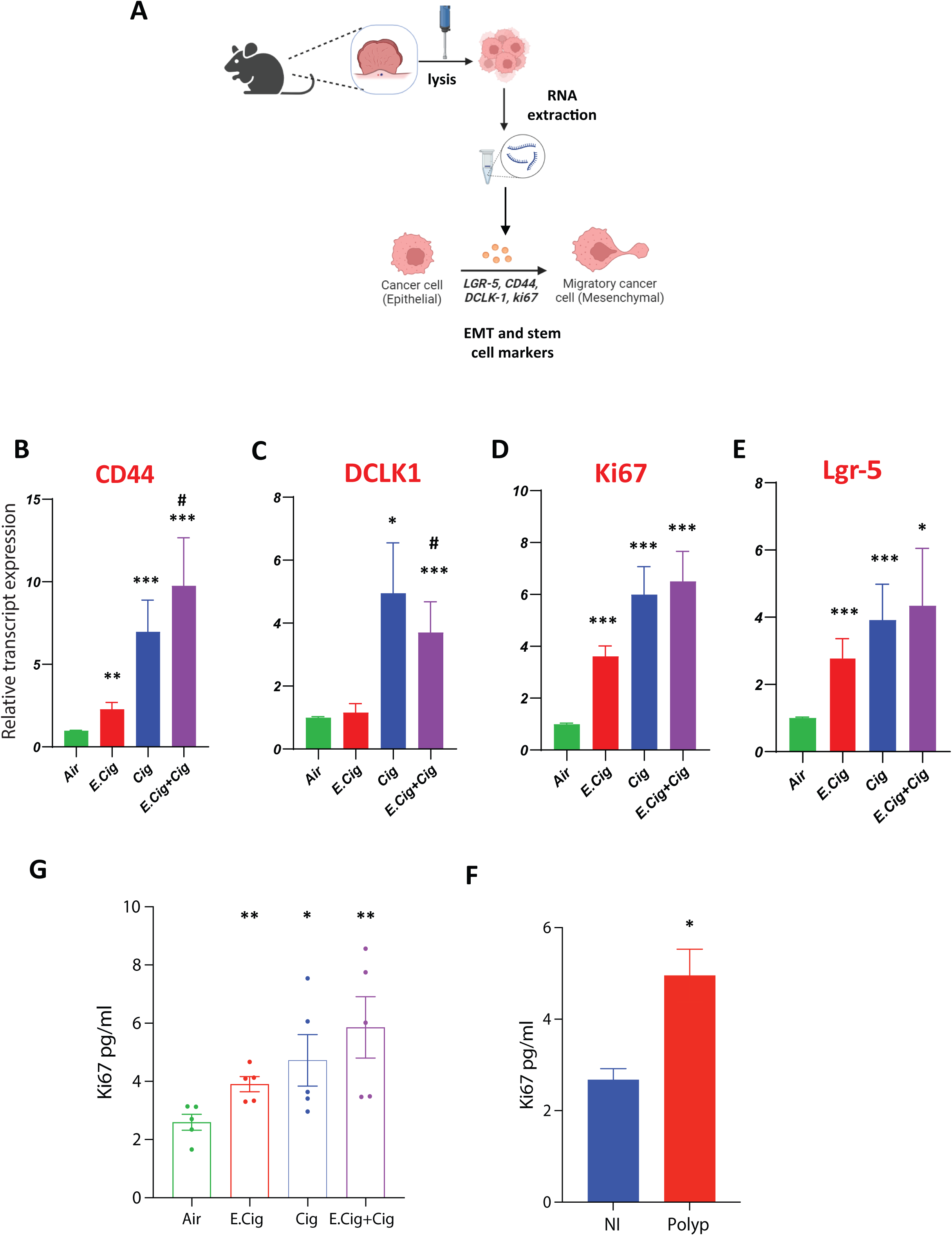
E.cig and Dual exposure upregulates stemness and proliferation markers in the colon. **(A)** The experimental design showing analysis of the mRNA levels of cancer stem cell markers and EMT from the polyp of each exposed group. **(B-E)**The total RNA extracted from the polyp of each exposed group was assessed for CD-44 (B), DCLK-1 (C), ki67 (D), and Lgr-5 (E) by RT-qPCR. (F) The level of Ki67 in the whole polyp extract was measured by ELISA (n=5 per each group). Each dot represents one mouse. Data presented as means ± SEM. \**P*-value < 0.05, \*\**P*-value < 0.01, and \*\*\**P*-value < 0.001, as determined by the Mann-Whitney test. * indicates comparison to air-exposed controls; # indicates comparison to E.cig exposed mice, P < 0.05.

## Discussion

The study aims to determine the impact of E.cig on the development of CRC. The role of tobacco and cigarette smoke in fueling CRCs has been established through robust epidemiologic studies with long-term follow-up and substantiated by mechanistic studies [15,16,24,38]. Since many young teenagers shifted from conventional smoking to E.cig or use both (dual), it is crucial to study the hazardous effects of E.cig and/or dual exposure on human health. E.cig vaping causes several respiratory complications including lung injury, lung edema, air epithelial pathway injury, and tissue hypoxia [57]. Besides, E.cig increases the risk of cardiovascular complications, induces endothelial dysfunction, and stimulates systemic inflammation [57]. On the gut, E.cig vaping mediates intestinal barrier defects by activating the inflammatory responses and affecting the gut epithelial tight junctions [10]. Although a leaky gut is a risk factor for many chronic diseases, including CRC, there is no report on the direct effect of E.cig on the development of CRC. Regarding dual exposure (E.cig +cig), a recent study showed that dual exposure had a worse outcome on the immune system and lung than the single exposure [58]

We used a well-known CPC-APC or CPC-APC^Min/+^murine model. APC^Min/+^ mice have a point mutation in the Apc gene and are a model for human FAP syndrome. Given the high prevalence of APC mutations in sporadic colorectal cancer, these models recapitulate most, if not all genetic and phenotypic aspects of CRC initiation and progression in humans [23,59]. The CPC-APC conditional knockout specifically develops adenomas/polyps in the colon/rectum, which are apparent around 3 months of age [23,59]. This model was used by other researchers to assess the impact of cigarette smoke on CRC progression and study the mechanistic pathways of cigarette smoke-mediate CRC [24]. Although there are other CRC mice models, however, these models were not used in studies related to E.cig or cig smoking.

In this study, we assessed the impact of E.cig vaping on CRC by several microscopic, molecular, and immunological approaches. Mice exposed to E.cig, cig, or both developed a significantly higher number of polyps and a higher polyp area compared to the air-exposed mice, suggesting that E.cig and cig can fuel CRC progression. Interestingly, mice exposed to both E.cig and cig (dual) had a significantly higher number of polyps compared to mice exposed to E.cig vapor alone, suggesting that administration of both E.cig and cig is more hazardous than vaping E.cig alone. Similarly, Hamon and colleagues showed that dual exposure had more damaging effect on lung epithelial cells and in the functions of phagocyte compared to the single exposure [58]

Next, we studied the possible pathways through which E.cig mediates CRC progression. We focused on inflammatory pathways, host DNA damage and repair pathways, and stemness markers associated with cancer stem cells. Chronic inflammation in the gut is fueling CRC, and patients with inflammatory bowel diseases are at increased risk of CRC development [40]. Our previous study showed that E.cig vaping stimulated the inflammatory signaling pathways in the gut of healthy subjects resulting in a leaky gut [10]. Herein, we found that E.cig, cig, and dual exposure stimulated the inflammatory signaling pathways in the polyp regions of the colon as shown by an increase in the infiltration of immune cells, pathology score, and upregulated the transcript levels of inflammatory cytokines such as IL-6, MCP-1, TNF-α, and Cxcl-1. The transcript levels were higher in dual-exposed mice than in E-cig-exposed mice. Likewise, using in vitro THP1 and monocyte differentiated macrophage cells from healthy non-smoker, dual exposure stimulated higher IL-8 than the single exposure [58]. These data suggest that E.cig could mediate CRC via activation of the inflammatory pathways.

Then, we assessed the impact of E.cig on host DNA damage and host DNA repair proteins, more specifically base excision repair pathways and DNA glycosylases. We focused on DNA glycosylases, especially NEIL-1 and NEIL-2 enzymes since our previous studies showed that gut pathogens can stimulate DNA damage and promote gastrointestinal cancers by suppression of NEIL-2 at both transcriptional and translational levels [33,34]. Our data showed that E.cig, cig, and dual exposures induce DNA breaks, and cause specific DNA double-strand breaks. Besides E.cig, cig, and dual exposures suppress DNA repair enzymes such as NEIL-2, NEIL-1, and NTH1, but not OGG1. The previous effects result in the accumulation of mutations and excessive DNA damage leading to genome instability and cancer. Similar to the inflammatory pathway, the effect of dual exposures on DNA damage was higher than E.cig exposure. A previous study by Lee and colleagues showed that aldehydes present in smoking and/or E.cig reduced the expression of nucleotide excision repair (NER) and base excision repair (BER) enzymes, more specifically proteins XPC (associated with NER) and 8-oxo guanine DNA glycosylase (OGG1/2, associated with BER) that are playing significant roles in assessing DNA damage mediated by O6-methyl-deoxuguanosine and initiating DNA repair [42]. The nicotine present in E.cig is metabolized to toxic metabolites such as N-nitrosonornicotine (NNN) and nicotine-derived nitrosamine ketones (NNK), the latter was further degraded to O6-methyl-deoxuguanosine that increased the mutagenesis in human lung and bladder epithelial cells [42]. The reduced expression or mutations of NEIL enzymes and OGG1 are associated with an increased risk of cancers [30,60–62]. In the study, E.cig vaping did not affect the level of OGG1 in the mice gut. The discrepancy between our data and the previous study could be due to the organ analyzed, further studies need to confirm this point. Collectively, these findings indicate that E.cig can promote CRC through suppression of host BER pathways and inducing double-strand DNA breaks resulting in the accumulation of mutations and genomic instability.

Here, we also investigated if E.cig vapor affects CSCs in the colon of exposed mice. CSCs play several roles in metastasis, angiogenesis, and EMT [49]. We focused on specific markers of CSCs such as CD44, Lgr-5, and DCLk1. CD44 is a surface glycoprotein, in which its expression is controlled by β-catenin/Tcf-4 signaling pathway. The overexpression of CD44 is associated with the loss of the APC gene and it is an early marker in adenoma-CRC progression [55]. CD44 and CD44v isoforms are markers of CSCs and targeting these molecules can be potential cancer therapeutic agents [63]. Deletion of CD44 from APC^Min/+^ mice reduces intestinal tumorigenesis [64]. Leucine-rich-repeat containing G-protein-coupled receptor 5 (Lgr5) is another biomarker of CSCs, and it is a receptor for WNT ligands such as R-spondin. LGR-5 positive cells act as CSCs and promote cancer propagation through clonal expansion [56]. The LGR-5 positive cells in adenoma can replicate producing more similar cells and other types of adenoma cell types [65]. Importantly, the Loss of the APC gene can stimulate CSCs and induce transformation in the LGR-5 positive cells to promote adenoma-to-carcinoma progression [66]. Doublecortin-like kinase 1 (DCLK1) is a potential label for CRC CSCs. It is overexpressed in CSCs and maintains the self-renewal and NOTCH1 survival signaling of CSCs resulting in increasing the tumor colony in APC^Min/+^ mice [52]. DCLK1 also regulates tumor microenvironment and immune cell infiltration in CRC [67]. Ki67 controls the transcriptome changes and steps of tumorigenesis. APC mice lacking Ki67 do not develop tumors [68]. We found that E.cig, cig, and dual exposures increase the gene expression of *CD44, Ki67, and Lgr-5* transcripts. Not only the transcript level, but they also significantly increased Ki67 protein levels compared to air-exposed mice. For *Dclk1*, cig and dual exposures upregulate the transcript level, but E.cig was not. It is not clear why E.cig alone could not upregulate the *Dclk1* level. Future studies need to verify this point. Collectively, our data showed that E.cig and other vaping exposures activate the markers of CSCs in the tumor regions to mediate tumor proliferation and expansion.

This study has some advantages and limitations. It is the first study that showed E.cig exposure could affect CRC progression and it shows possible pathways responsible for this progression such as activation of inflammatory signaling, inducing DNA damage and suppression of DNA repair proteins, and activation of cancer stem cells. The limitation of the study is the number of CPC-APC mice used in cig and dual exposure is not large enough. Future studies are required including a) a bigger cohort of the mice with different time points, b) other models of CRC mice and c) other brands of cig and e-cig. However, our results consistently revealed that dual exposure poses a higher risk compared to single E-cig exposure. Here we showed a selected number of pathways associated with CRC following exposure. Future studies are ongoing to understand the global pathways that link E.cig exposures with CRC development.

In conclusion, we reported here that E.cig could promote CRC in CPC-APC mice by stimulating inflammatory pathways, activation of DNA damage, and promoting the activity of cancer stem cells. It is crucial to alert the public about the possible hazardous effects of E.cig alone or in combination with conventional smoking.

## Author Contribution

IMS, SD and LCA design the study. IMS, KI, LD, ID, and KP performed animal experiments; AC performed western blot and LA-PCR assay; ST: helped in IF imaging, IMS, AC, HTK, SD, and LCA analyzed data; HKT for resources of DNA damage assays, SD and LCA supervised the research. IMS wrote the original draft, SD, LCA edited and revised the manuscript. All authors edited and agreed on the final version of the manuscript.

## Acknowledgment

We are grateful to UMass-Lowell CoRE Research Facility (CRF) Cell Analysis & Imaging Lab for the confocal microscopy used in this study. We thank the histology core facility at UMASS Chan Medical School (Histospring.com) for staining γH2AX in tissue specimens. We also thank Joshua Alcantara for technical assistance in mice breeding and euthanasia. Diagrams were designed using free software https://www.biorender.com/.

## Data availability

The data that support the findings of this study are present in the main text. For further inquiries, please contact the corresponding authors.

## Ethics approval

The manuscript includes animal experiments. The experimental design was approved by the University of California San Diego Institutional Animal Care and Use Committee (IACUC) policies under the animal protocol number (S18086 and S16021). No human data in this study.

## References

1. McCarthy, M. E-cigarettes are major threat to young people’s health, says US surgeon general. 2016.

2. Mirbolouk, M.; Charkhchi, P.; Kianoush, S.; Uddin, S.I.; Orimoloye, O.A.; Jaber, R.; Bhatnagar, A.; Benjamin, E.J.; Hall, M.E.; DeFilippis, A.P. Prevalence and distribution of e-cigarette use among US adults: behavioral risk factor surveillance system, 2016. Annals of internal medicine 2018, 169, 429–438.

3. Regan, A.K.; Promoff, G.; Dube, S.R.; Arrazola, R. Electronic nicotine delivery systems: adult use and awareness of the ‘e-cigarette’ in the USA. Tob Control 2013, 22, 19–23, doi:10.1136/tobaccocontrol-2011-050044.

4. Gentzke, A.S.; Creamer, M.; Cullen, K.A.; Ambrose, B.K.; Willis, G.; Jamal, A.; King, B.A. Vital signs: tobacco product use among middle and high school students—United States, 2011–2018. Morbidity and Mortality Weekly Report 2019, 68, 157.

5. Zhu, S.-H.; Sun, J.Y.; Bonnevie, E.; Cummins, S.E.; Gamst, A.; Yin, L.; Lee, M. Four hundred and sixty brands of e-cigarettes and counting: implications for product regulation. Tobacco control 2014, 23, iii3–iii9.

6. O’Loughlin, J.; Wellman, R.J.; Potvin, L. Whither the e-cigarette? Int J Public Health 2016, 61, 147–148, doi:10.1007/s00038-016-0800-5.

7. King, B.A.; Jones, C.M.; Baldwin, G.T.; Briss, P.A. The EVALI and Youth Vaping Epidemics - Implications for Public Health. N Engl J Med 2020, 382, 689–691, doi:10.1056/NEJMp1916171.

8. Shinbashi, M.; Rubin, B.K. Electronic cigarettes and e-cigarette/vaping product use associated lung injury (EVALI). Paediatr Respir Rev 2020, 36, 87–91, doi:10.1016/j.prrv.2020.06.003.

9. Sayed, I.M.; Masso-Silva, J.A.; Mittal, A.; Patel, A.; Lin, E.; Moshensky, A.; Shin, J.; Bojanowski, C.M.; Das, S.; Akuthota, P.;, et al. Inflammatory phenotype modulation in the respiratory tract and systemic circulation of e-cigarette users: a pilot study. Am J Physiol Lung Cell Mol Physiol 2021, 321, L1134–l1146, doi:10.1152/ajplung.00363.2021.

10. Sharma, A.; Lee, J.; Fonseca, A.G.; Moshensky, A.; Kothari, T.; Sayed, I.M.; Ibeawuchi, S.R.; Pranadinata, R.F.; Ear, J.; Sahoo, D.;, et al. E-cigarettes compromise the gut barrier and trigger inflammation. iScience 2021, 24, 102035, doi:10.1016/j.isci.2021.102035.

11. Genua, F.; Raghunathan, V.; Jenab, M.; Gallagher, W.M.; Hughes, D.J. The Role of Gut Barrier Dysfunction and Microbiome Dysbiosis in Colorectal Cancer Development. Front Oncol 2021, 11, 626349, doi:10.3389/fonc.2021.626349.

12. Diergaarde, B.; Vrieling, A.; van Kraats, A.A.; van Muijen, G.N.; Kok, F.J.; Kampman, E. Cigarette smoking and genetic alterations in sporadic colon carcinomas. Carcinogenesis 2003, 24, 565–571, doi:10.1093/carcin/24.3.565.

13. Sarebø, M.; Skjelbred, C.F.; Breistein, R.; Lothe, I.M.; Hagen, P.C.; Bock, G.; Hansteen, I.L.; Kure, E.H. Association between cigarette smoking, APC mutations and the risk of developing sporadic colorectal adenomas and carcinomas. BMC Cancer 2006, 6, 71, doi:10.1186/1471-2407-6-71.

14. Lüchtenborg, M.; Weijenberg, M.P.; Kampman, E.; van Muijen, G.N.; Roemen, G.M.; Zeegers, M.P.; Goldbohm, R.A.; van ’t Veer, P.; de Goeij, A.F.; van den Brandt, P.A. Cigarette smoking and colorectal cancer: APC mutations, hMLH1 expression, and GSTM1 and GSTT1 polymorphisms. Am J Epidemiol 2005, 161, 806–815, doi:10.1093/aje/kwi114.

15. Liang, P.S.; Chen, T.Y.; Giovannucci, E. Cigarette smoking and colorectal cancer incidence and mortality: systematic review and meta-analysis. Int J Cancer 2009, 124, 2406–2415, doi:10.1002/ijc.24191.

16. Botteri, E.; Borroni, E.; Sloan, E.K.; Bagnardi, V.; Bosetti, C.; Peveri, G.; Santucci, C.; Specchia, C.; van den Brandt, P.; Gallus, S.;, et al. Smoking and Colorectal Cancer Risk, Overall and by Molecular Subtypes: A Meta-Analysis. Am J Gastroenterol 2020, 115, 1940–1949, doi:10.14309/ajg.0000000000000803.

17. Tsoi, K.K.; Pau, C.Y.; Wu, W.K.; Chan, F.K.; Griffiths, S.; Sung, J.J. Cigarette smoking and the risk of colorectal cancer: a meta-analysis of prospective cohort studies. Clin Gastroenterol Hepatol 2009, 7, 682–688.e681-685, doi:10.1016/j.cgh.2009.02.016.

18. Siegel, R.L.; Wagle, N.S.; Cercek, A.; Smith, R.A.; Jemal, A. Colorectal cancer statistics, 2023. CA Cancer J Clin 2023, 73, 233–254, doi:10.3322/caac.21772.

19. Sayed, I.M.; El-Hafeez, A.A.A.; Maity, P.P.; Das, S.; Ghosh, P. Modeling colorectal cancers using multidimensional organoids. Adv Cancer Res 2021, 151, 345–383, doi:10.1016/bs.acr.2021.02.005.

20. Morin, P.J.; Sparks, A.B.; Korinek, V.; Barker, N.; Clevers, H.; Vogelstein, B.; Kinzler, K.W. Activation of beta-catenin-Tcf signaling in colon cancer by mutations in beta-catenin or APC. Science 1997, 275, 1787–1790, doi:10.1126/science.275.5307.1787.

21. Zhang, L.; Shay, J.W. Multiple Roles of APC and its Therapeutic Implications in Colorectal Cancer. J Natl Cancer Inst 2017, 109, doi:10.1093/jnci/djw332.

22. Groden, J.; Thliveris, A.; Samowitz, W.; Carlson, M.; Gelbert, L.; Albertsen, H.; Joslyn, G.; Stevens, J.; Spirio, L.; Robertson, M.;, et al. Identification and characterization of the familial adenomatous polyposis coli gene. Cell 1991, 66, 589–600, doi:10.1016/0092-8674(81)90021-0.

23. Hinoi, T.; Akyol, A.; Theisen, B.K.; Ferguson, D.O.; Greenson, J.K.; Williams, B.O.; Cho, K.R.; Fearon, E.R. Mouse model of colonic adenoma-carcinoma progression based on somatic Apc inactivation. Cancer Res 2007, 67, 9721–9730, doi:10.1158/0008-5472.Can-07-2735.

24. Wang, H.; Chen, X.; Gao, Q.; Liu, K.; Bi, G.; Deng, J.; Zhang, X. Smoking induces the occurrence of colorectal cancer via changing the intestinal permeability. J buon 2021, 26, 1009–1015.

25. Salamanca, J.C.; Meehan-Atrash, J.; Vreeke, S.; Escobedo, J.O.; Peyton, D.H.; Strongin, R.M. E-cigarettes can emit formaldehyde at high levels under conditions that have been reported to be non-averse to users. Scientific reports 2018, 8, 1–6.

26. Perez, M.F.; Crotty Alexander, L.E. Why is vaping going up in flames? Annals of the American Thoracic Society 2020, 17, 545–549.

27. Bozier, J.; Chivers, E.K.; Chapman, D.G.; Larcombe, A.N.; Bastian, N.; Masso-Silva, J.A.; Byun, M.K.; McDonald, C.F.; Alexander Crotty, L.E.; Ween, M.P. The Evolving Landscape of Electronic Cigarettes: A systematic review of recent evidence. Chest 2020, doi:10.1016/j.chest.2019.12.042.

28. Ganapathy, V.; Manyanga, J.; Brame, L.; McGuire, D.; Sadhasivam, B.; Floyd, E.; Rubenstein, D.A.; Ramachandran, I.; Wagener, T.; Queimado, L. Electronic cigarette aerosols suppress cellular antioxidant defenses and induce significant oxidative DNA damage. PLoS One 2017, 12, e0177780, doi:10.1371/journal.pone.0177780.

29. Yu, V.; Rahimy, M.; Korrapati, A.; Xuan, Y.; Zou, A.E.; Krishnan, A.R.; Tsui, T.; Aguilera, J.A.; Advani, S.; Crotty Alexander, L.E.;, et al. Electronic cigarettes induce DNA strand breaks and cell death independently of nicotine in cell lines. Oral Oncol 2016, 52, 58–65, doi:10.1016/j.oraloncology.2015.10.018.

30. Krokan, H.E.; Bjørås, M. Base excision repair. Cold Spring Harb Perspect Biol 2013, 5, a012583, doi:10.1101/cshperspect.a012583.

31. Meira, L.B.; Bugni, J.M.; Green, S.L.; Lee, C.W.; Pang, B.; Borenshtein, D.; Rickman, B.H.; Rogers, A.B.; Moroski-Erkul, C.A.; McFaline, J.L.;, et al. DNA damage induced by chronic inflammation contributes to colon carcinogenesis in mice. J Clin Invest 2008, 118, 2516–2525, doi:10.1172/jci35073.

32. Dey, S.; Maiti, A.K.; Hegde, M.L.; Hegde, P.M.; Boldogh, I.; Sarkar, P.S.; Abdel-Rahman, S.Z.; Sarker, A.H.; Hang, B.; Xie, J.;, et al. Increased risk of lung cancer associated with a functionally impaired polymorphic variant of the human DNA glycosylase NEIL2. DNA Repair (Amst) 2012, 11, 570–578, doi:10.1016/j.dnarep.2012.03.005.

33. Sayed, I.M.; Chakraborty, A.; Abd El-Hafeez, A.A.; Sharma, A.; Sahan, A.Z.; Huang, W.J.M.; Sahoo, D.; Ghosh, P.; Hazra, T.K.; Das, S. The DNA Glycosylase NEIL2 Suppresses Fusobacterium-Infection-Induced Inflammation and DNA Damage in Colonic Epithelial Cells. Cells 2020, 9, doi:10.3390/cells9091980.

34. Sayed, I.M.; Sahan, A.Z.; Venkova, T.; Chakraborty, A.; Mukhopadhyay, D.; Bimczok, D.; Beswick, E.J.; Reyes, V.E.; Pinchuk, I.; Sahoo, D.;, et al. Helicobacter pylori infection downregulates the DNA glycosylase NEIL2, resulting in increased genome damage and inflammation in gastric epithelial cells. J Biol Chem 2020, 295, 11082–11098, doi:10.1074/jbc.RA119.009981.

35. Erben, U.; Loddenkemper, C.; Doerfel, K.; Spieckermann, S.; Haller, D.; Heimesaat, M.M.; Zeitz, M.; Siegmund, B.; Kühl, A.A. A guide to histomorphological evaluation of intestinal inflammation in mouse models. Int J Clin Exp Pathol 2014, 7, 4557–4576.

36. Schmittgen, T.D.; Livak, K.J. Analyzing real-time PCR data by the comparative C(T) method. Nat Protoc 2008, 3, 1101–1108, doi:10.1038/nprot.2008.73.

37. Chakraborty, A.; Wakamiya, M.; Venkova-Canova, T.; Pandita, R.K.; Aguilera-Aguirre, L.; Sarker, A.H.; Singh, D.K.; Hosoki, K.; Wood, T.G.; Sharma, G.;, et al. Neil2-null Mice Accumulate Oxidized DNA Bases in the Transcriptionally Active Sequences of the Genome and Are Susceptible to Innate Inflammation. J Biol Chem 2015, 290, 24636–24648, doi:10.1074/jbc.M115.658146.

38. Bai, X.; Wei, H.; Liu, W.; Coker, O.O.; Gou, H.; Liu, C.; Zhao, L.; Li, C.; Zhou, Y.; Wang, G.;, et al. Cigarette smoke promotes colorectal cancer through modulation of gut microbiota and related metabolites. Gut 2022, 71, 2439–2450, doi:10.1136/gutjnl-2021-325021.

39. Zhou, R.W.; Harpaz, N.; Itzkowitz, S.H.; Parsons, R.E. Molecular mechanisms in colitis-associated colorectal cancer. Oncogenesis 2023, 12, 48, doi:10.1038/s41389-023-00492-0.

40. Nardone, O.M.; Zammarchi, I.; Santacroce, G.; Ghosh, S.; Iacucci, M. Inflammation-Driven Colorectal Cancer Associated with Colitis: From Pathogenesis to Changing Therapy. Cancers (Basel) 2023, 15, doi:10.3390/cancers15082389.

41. Hanahan, D.; Weinberg, R.A. Hallmarks of cancer: the next generation. Cell 2011, 144, 646–674, doi:10.1016/j.cell.2011.02.013.

42. Lee, H.W.; Park, S.H.; Weng, M.W.; Wang, H.T.; Huang, W.C.; Lepor, H.; Wu, X.R.; Chen, L.C.; Tang, M.S. E-cigarette smoke damages DNA and reduces repair activity in mouse lung, heart, and bladder as well as in human lung and bladder cells. Proc Natl Acad Sci U S A 2018, 115, E1560–e1569, doi:10.1073/pnas.1718185115.

43. Hirata, N.; Horinouchi, T.; Kanda, Y. Effects of cigarette smoke extract derived from heated tobacco products on the proliferation of lung cancer stem cells. Toxicol Rep 2022, 9, 1273–1280, doi:10.1016/j.toxrep.2022.06.001.

44. Qian, W.; Kong, X.; Zhang, T.; Wang, D.; Song, J.; Li, Y.; Li, X.; Geng, H.; Min, J.; Kong, Q.;, et al. Cigarette smoke stimulates the stemness of renal cancer stem cells via Sonic Hedgehog pathway. Oncogenesis 2018, 7, 24, doi:10.1038/s41389-018-0029-7.

45. Sun, X.; Song, J.; Li, E.; Geng, H.; Li, Y.; Yu, D.; Zhong, C. Cigarette smoke supports stemness and epithelial-mesenchymal transition in bladder cancer stem cells through SHH signaling. Int J Clin Exp Pathol 2020, 13, 1333–1348.

46. Lin, S.; Mei, W.; Lai, H.; Li, X.; Weng, H.; Xiong, J.; Lin, X.; Zeng, T.; Zhang, Q.; Liu, X.;, et al. Cigarette smoking promotes keratinocyte malignancy via generation of cancer stem-like cells. Journal of Cancer 2021, 12, 1085–1093, doi:10.7150/jca.50746.

47. Xie, C.; Zhu, J.; Wang, X.; Chen, J.; Geng, S.; Wu, J.; Zhong, C.; Li, X. Tobacco smoke induced hepatic cancer stem cell-like properties through IL-33/p38 pathway. Journal of Experimental & Clinical Cancer Research 2019, 38, 39, doi:10.1186/s13046-019-1052-z.

48. Nimmakayala, R.K.; Seshacharyulu, P.; Lakshmanan, I.; Rachagani, S.; Chugh, S.; Karmakar, S.; Rauth, S.; Vengoji, R.; Atri, P.; Talmon, G.A.;, et al. Cigarette Smoke Induces Stem Cell Features of Pancreatic Cancer Cells via PAF1. Gastroenterology 2018, 155, 892–908.e896, 10.1053/j.gastro.2018.05.041.

49. Liu, Q.; Guo, Z.; Li, G.; Zhang, Y.; Liu, X.; Li, B.; Wang, J.; Li, X. Cancer stem cells and their niche in cancer progression and therapy. Cancer Cell International 2023, 23, 305, doi:10.1186/s12935-023-03130-2.

50. Thapa, R.; Wilson, G.D. The Importance of CD44 as a Stem Cell Biomarker and Therapeutic Target in Cancer. Stem Cells Int 2016, 2016, 2087204, doi:10.1155/2016/2087204.

51. Ihemelandu, C.; Naeem, A.; Parasido, E.; Berry, D.; Chaldekas, K.; Harris, B.T.; Rodriguez, O.; Albanese, C. Clinicopathologic and prognostic significance of LGR5, a cancer stem cell marker in patients with colorectal cancer. Colorectal Cancer 2019, 8, Crc11, doi:10.2217/crc-2019-0009.

52. Chandrakesan, P.; Yao, J.; Qu, D.; May, R.; Weygant, N.; Ge, Y.; Ali, N.; Sureban, S.M.; Gude, M.; Vega, K.;, et al. Dclk1, a tumor stem cell marker, regulates pro-survival signaling and self-renewal of intestinal tumor cells. Molecular Cancer 2017, 16, 30, doi:10.1186/s12943-017-0594-y.

53. Cidado, J.; Wong, H.Y.; Rosen, D.M.; Cimino-Mathews, A.; Garay, J.P.; Fessler, A.G.; Rasheed, Z.A.; Hicks, J.; Cochran, R.L.; Croessmann, S.;, et al. Ki-67 is required for maintenance of cancer stem cells but not cell proliferation. Oncotarget 2016, 7, 6281–6293, doi:10.18632/oncotarget.7057.

54. Abbasi, N.; Long, T.; Li, Y.; Yee, B.A.; Cho, B.S.; Hernandez, J.E.; Ma, E.; Patel, P.R.; Sahoo, D.; Sayed, I.M.;, et al. DDX5 promotes oncogene C3 and FABP1 expressions and drives intestinal inflammation and tumorigenesis. Life Sci Alliance 2020, 3, doi:10.26508/lsa.202000772.

55. Wielenga, V.J.; Smits, R.; Korinek, V.; Smit, L.; Kielman, M.; Fodde, R.; Clevers, H.; Pals, S.T. Expression of CD44 in Apc and Tcf mutant mice implies regulation by the WNT pathway. Am J Pathol 1999, 154, 515–523, doi:10.1016/s0002-9440(10)65297-2.

56. Yanai, H.; Atsumi, N.; Tanaka, T.; Nakamura, N.; Komai, Y.; Omachi, T.; Tanaka, K.; Ishigaki, K.; Saiga, K.; Ohsugi, H.;, et al. Intestinal cancer stem cells marked by Bmi1 or Lgr5 expression contribute to tumor propagation via clonal expansion. Sci Rep 2017, 7, 41838, doi:10.1038/srep41838.

57. Marques, P.; Piqueras, L.; Sanz, M.-J. An updated overview of e-cigarette impact on human health. Respiratory Research 2021, 22, 151, doi:10.1186/s12931-021-01737-5.

58. Hamon, R.; Thredgold, L.; Wijenayaka, A.; Bastian, N.A.; Ween, M.P. Dual Exposure to E-Cigarette Vapour and Cigarette Smoke Results in Poorer Airway Cell, Monocyte, and Macrophage Function Than Single Exposure. Int J Mol Sci 2024, 25, doi:10.3390/ijms25116071.

59. Drummond, F.; Sowden, J.; Morrison, K.; Edwards, Y.H. The caudal-type homeobox protein Cdx-2 binds to the colon promoter of the carbonic anhydrase 1 gene. Eur J Biochem 1996, 236, 670–681, doi:10.1111/j.1432-1033.1996.t01-1-00670.x.

60. Shinmura, K.; Kato, H.; Kawanishi, Y.; Igarashi, H.; Goto, M.; Tao, H.; Inoue, Y.; Nakamura, S.; Misawa, K.; Mineta, H.;, et al. Abnormal Expressions of DNA Glycosylase Genes NEIL1, NEIL2, and NEIL3 Are Associated with Somatic Mutation Loads in Human Cancer. Oxid Med Cell Longev 2016, 2016, 1546392, doi:10.1155/2016/1546392.

61. Zhai, X.; Zhao, H.; Liu, Z.; Wang, L.E.; El-Naggar, A.K.; Sturgis, E.M.; Wei, Q. Functional variants of the NEIL1 and NEIL2 genes and risk and progression of squamous cell carcinoma of the oral cavity and oropharynx. Clin Cancer Res 2008, 14, 4345–4352, doi:10.1158/1078-0432.Ccr-07-5282.

62. Ali, K.; Mahjabeen, I.; Sabir, M.; Mehmood, H.; Kayani, M.A. OGG1 Mutations and Risk of Female Breast Cancer: Meta-Analysis and Experimental Data. Disease Markers 2015, 2015, 690878, doi:10.1155/2015/690878.

63. Yan, Y.; Zuo, X.; Wei, D. Concise Review: Emerging Role of CD44 in Cancer Stem Cells: A Promising Biomarker and Therapeutic Target. Stem Cells Transl Med 2015, 4, 1033–1043, doi:10.5966/sctm.2015-0048.

64. Zeilstra, J.; Joosten, S.P.; Dokter, M.; Verwiel, E.; Spaargaren, M.; Pals, S.T. Deletion of the WNT target and cancer stem cell marker CD44 in Apc(Min/+) mice attenuates intestinal tumorigenesis. Cancer Res 2008, 68, 3655–3661, doi:10.1158/0008-5472.Can-07-2940.

65. Schepers, A.G.; Snippert, H.J.; Stange, D.E.; van den Born, M.; van Es, J.H.; van de Wetering, M.; Clevers, H. Lineage tracing reveals Lgr5+ stem cell activity in mouse intestinal adenomas. Science 2012, 337, 730–735, doi:10.1126/science.1224676.

66. Fu, T.; Coulter, S.; Yoshihara, E.; Oh, T.G.; Fang, S.; Cayabyab, F.; Zhu, Q.; Zhang, T.; Leblanc, M.; Liu, S.;, et al. FXR Regulates Intestinal Cancer Stem Cell Proliferation. Cell 2019, 176, 1098–1112.e1018, doi:10.1016/j.cell.2019.01.036.

67. Wu, X.; Qu, D.; Weygant, N.; Peng, J.; Houchen, C.W. Cancer Stem Cell Marker DCLK1 Correlates with Tumorigenic Immune Infiltrates in the Colon and Gastric Adenocarcinoma Microenvironments. Cancers (Basel) 2020, 12, doi:10.3390/cancers12020274.

68. Mrouj, K.; Andrés-Sánchez, N.; Dubra, G.; Singh, P.; Sobecki, M.; Chahar, D.; Al Ghoul, E.; Aznar, A.B.; Prieto, S.; Pirot, N.;, et al. Ki-67 regulates global gene expression and promotes sequential stages of carcinogenesis. Proc Natl Acad Sci U S A 2021, 118, doi:10.1073/pnas.2026507118.

